# Fact retrieval or compacted procedures in arithmetic – a neurophysiological investigation of two hypotheses

**DOI:** 10.1101/2020.03.10.985143

**Authors:** Roland H. Grabner, Clemens Brunner, Valerie Lorenz, Stephan E. Vogel, Bert De Smedt

**Affiliations:** Institute of Psychology, University of Graz, Universitätsplatz 2, 8010 Graz, Austria; Parenting and Special Education Research Unit, KU Leuven, Leopold Vanderkelenstraat 32 – bus 3765, 3000 Leuven, Belgium

**Keywords:** arithmetic, problem-solving strategies, induced EEG, ERS/ERD, fact retrieval, compacted procedures

## Abstract

There is broad consensus that adults solve single-digit multiplication problems almost exclusively by fact retrieval (i.e., retrieval of the solution from an arithmetic fact network). In contrast, there has been a long-standing debate on the cognitive processes involved in solving single-digit addition problems. This debate has evolved around two theoretical accounts. The *fact-retrieval account* postulates that these are solved through fact retrieval, just like multiplications, whereas the *compacted-procedure account* proposes that solving very small additions (i.e., problems with operands between 1 and 4) involves highly automatized and unconscious compacted procedures. In the present electroencephalography (EEG) study, we put these two accounts to the test by comparing neurophysiological correlates of solving very small additions and multiplications. A sample of 40 adults worked on an arithmetic production task involving all (non-tie) single-digit additions and multiplications. Afterwards, participants completed trial-by-trial strategy self-reports. In our EEG analyses, we focused on induced activity (event-related synchronization/desynchronization, ERS/ERD) in three frequency bands (theta, lower alpha, upper alpha). Across all frequency bands, we found higher evidential strength for similar rather than different neurophysiological processes accompanying the solution of very small addition and multiplication problems. This was also true when *n* + 1 and *n* × 1 problems were excluded from the analyses. In two additional analyses, we showed that ERS/ERD can differentiate between self-reported problem-solving strategies (retrieval vs. procedure) and even between *n* + 1 and *n* + *m* problems in very small additions, demonstrating its high sensitivity to cognitive processes in arithmetic. The present findings clearly support the fact-retrieval account, suggesting that both very small additions and multiplications are solved through fact retrieval.

**HIGHLIGHTS:**

- Neurophysiological test of fact retrieval and compacted procedures account
- Induced EEG data are sensitive to cognitive processes in arithmetic problem solving
- Both very small additions and multiplications are solved through fact retrieval

## 1. INTRODUCTION

After more than three decades of research on cognitive processes in arithmetic, there is broad consensus that fact retrieval (i.e., direct retrieval of the solution from an arithmetic fact network in long-term memory) is the dominant process for solving single-digit *multiplication* problems (e.g., 2 × 3) in adults (Ashcraft, 1992; Campbell & Epp, 2005). This view is in line with the observation that the multiplication table is verbally learned by rote in school and with results from several behavioral and neurophysiological studies. For instance, adults solve single-digit multiplication problems quickly (e.g., Groen & Parkman, 1972), usually indicate the use of fact retrieval in strategy reports (e.g., Campbell & Xue, 2001), and display brain activation patterns related to language processing (e.g., Prado et al., 2011). In contrast, there has been a long-standing debate on the cognitive processes involved in solving single-digit *addition* problems (e.g., 2 + 3), which are typically solved equally fast and for which individuals likewise report fact retrieval as their main solution strategy (Campbell & Xue, 2001). This debate has evolved around two major theoretical accounts: the *fact-retrieval account* and the *compacted-procedure account* (Baroody, 2018; Chen & Campbell, 2018). In the current electroencephalography (EEG) study, we used for the first time neurophysiological measures to pit these two accounts against each other and to shed new light on this long-standing debate.

The *fact-retrieval account*, on the one hand, proposes that educated adults solve single-digit addition and multiplication problems through a similar cognitive process, i.e., fact retrieval from long-term memory (e.g., Groen & Parkman, 1972; LeFevre, Sadesky, & Bisanz, 1996; McCloskey, Harley, & Sokol, 1991). Even though additions are usually not learned by rote as in multiplication, it is assumed that after enough practice in solving these problems they are eventually stored as arithmetic facts in long-term memory. Therefore, the solution to single-digit additions can also be rapidly retrieved.

The *compacted-procedure account*, on the other hand, postulates that some single-digit additions, in particular very small ones, are solved through a rapid and unconscious arithmetic procedure, also the result of years of practice (e.g., Barrouillet & Thevenot, 2013; Uittenhove, Thevenot, & Barrouillet, 2016). This procedure is assumed to be highly automatic (compacted) such that it is equally fast as fact retrieval and does not reach consciousness. Given the rapid problem-solving process, individuals would have the feeling of having retrieved a fact from memory and therefore would report this strategy.

The compacted-procedure account goes back to Baroody (1983) and has seen a revival in the past years due to behavioral findings that seem to be incompatible with the fact-retrieval account (e.g., Barrouillet & Thevenot, 2013; Uittenhove et al., 2016). These findings mainly come from detailed analyses of the problem-size effect in single-digit additions (Barrouillet & Thevenot, 2013; Uittenhove et al., 2016) and operator priming effects in multiplications and additions (Fayol & Thevenot, 2012; Roussel, Fayol, & Barrouillet, 2002).

The problem-size effect is a robust and well-established behavioral finding in arithmetic (for a review, cf. Zbrodoff & Logan, 2005), which is reflected in longer reaction times and lower accuracies when solving problems with larger operands (e.g., additions with sums > 10) compared to those with smaller operands (e.g., additions with sums ≤ 10). Barrouillet and Thevenot (2013) conducted a detailed investigation of the problem-size effect in single-digit additions. They presented addition problems with operands from 1 to 4 to adult students (*n =* 92) and found a clear problem-size effect in the non-tie problems (i.e., 12 problems with different operands): reaction times were highly correlated with the operands’ sum. The authors argued that a problem-size effect in these very small problems does not fit well with the fact-retrieval account, because retrieval times should not vary significantly between these overlearned problems, nor should they be strongly correlated with the operands’ magnitudes. Rather, these findings would be more in line with the hypothesis of “a compacted and automated procedure” (p. 44) that scrolls “an ordered representation such as a number line or a verbal number sequence” (p. 35). This scrolling would resemble a counting strategy, moving first to the position of the first operand and then counting on by the second operand to the correct sum.

Uittenhove et al. (2016) extended the study by Barrouillet and Thevenot (2013) by including the full set of single-digit addition problems and individual strategy reports. Specifically, 90 adults worked on all possible 81 single-digit addition problems (ties and non-ties) in two sessions. In the first session, all problems were presented six times; in the second session, after solving them again (once), participants reported the applied strategy for each problem. The self-reported frequency of fact retrieval was the best predictor of the reaction times in the entire set of non-tie problems. In the small non-tie problems (sums ≤ 10), however, reaction times displayed the highest correlations with the operands’ minimum and product. A plot of the sum-related reaction time increase (Uittenhove et al., 2016, Fig. 1) suggested that a problem-size effect emerged only for the smallest problems (sums between 3 and 7), whereas in the other small problems (sums between 7 and 10) no reaction time increase was observed. This pattern of findings led Uittenhove et al. to suggest that fact retrieval could not have been used for the smallest problems (i.e., with operands between 1 and 4). In an additional analysis, which is most relevant to the present study, they focused on a sub-sample of 51 participants who reported fact retrieval in all of the problems with operands between 1 and 4. Within these 12 very small problems, reaction time was highly correlated with the operands’ sum, as was also observed by Barrouillet and Thevenot (2013). In the medium small problems (sums from 7 to 10, without *n* + 1 problems; six problems), in contrast, no problem size effect was found. A similar finding emerged when the *n* + 1 problems were analyzed separately from the other problems. In the very small problems, both the six *n* + 1 and six *n* + *m* problems showed a significant problem size effect, which was stronger than that for the two problem categories in the medium small additions. Together, the findings in the very small problems were interpreted to be incompatible with the fact-retrieval account. Instead, the authors argued that the solutions to these addition problems are “reconstructed through a rapid sequential procedure the duration of which is determined by the magnitude of the operands”, and that “this rapid procedure seems to be limited in its application to very small operands that do not exceed 4” (p. 298). The medium small problems were assumed to be solved through a mixture of fact retrieval (in the *n* + *m* problems) or the application of a one-greater rule (in the *n* + 1 problems).

**Figure 1.**
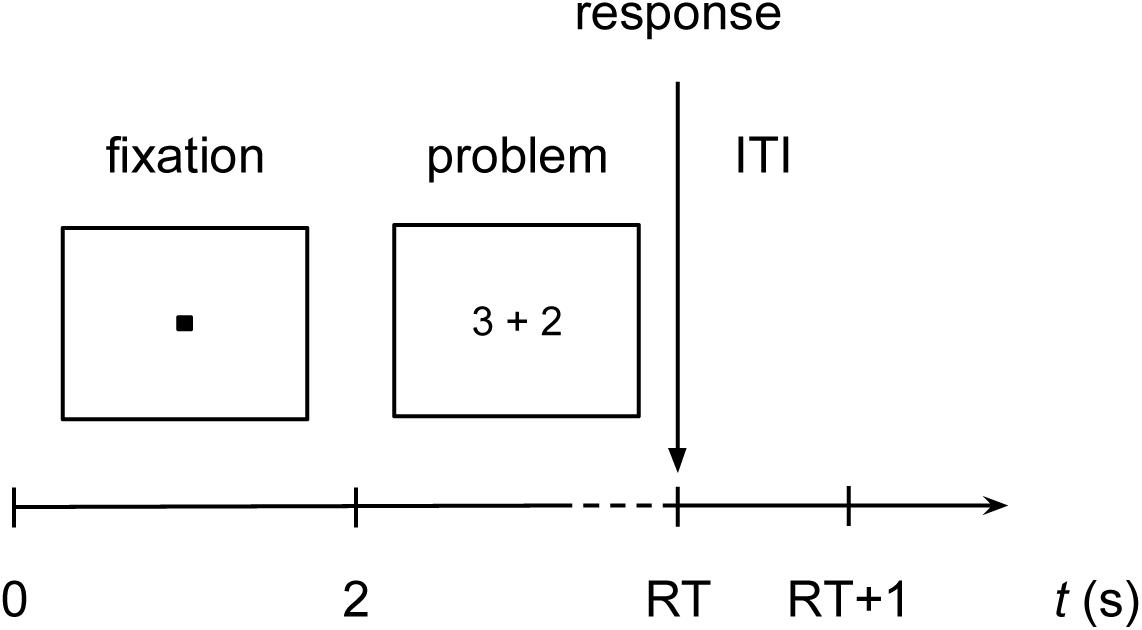
Schematic illustration of an EEG trial. RT indicates the reaction time from problem onset to oral response.

Further behavioral evidence for the compacted-procedure account comes from priming studies, in which multiplications and additions were compared. Roussel, Fayol, and Barrouillet (2002) as well as Fayol and Thevenot (2012) administered an operator priming task, in which the operation sign (i.e., “+” or “×”) was displayed briefly (e.g., 150 ms) before both arithmetic operands. This priming resulted in faster responses for additions and subtractions but not for multiplications. Interestingly, the priming effect in addition was independent of problem size and emerged for small (operands ≤ 5) as well as large problems (operands ≥ 5). The operation-dependent facilitation effect was attributed to a pre-activation of an arithmetic procedure which is later used to solve the problem. Because this pre-activation did not emerge for multiplications, it was again concluded that in single-digit additions, even in small problems, procedural processes take place.

The conclusion of compacted procedures in single-digit additions based on behavioral findings from these and other studies (Mathieu, Gourjon, Couderc, Thevenot, & Prado, 2016; Thevenot, Barrouillet, Castel, & Uittenhove, 2016; Thevenot, Fanget, & Fayol, 2007), however, has not been met without criticism. Chen and Campbell (2018) recently reviewed the current evidence and concluded that the idea of compacted procedures “is not convincing and does not justify significant revision of the long-standing assumption in cognitive science that direct memory retrieval is ultimately the most efficient process of simple addition” (p. 751; but see also Baroody, 2018). Based on re-analyses of previous data, Chen and Campbell argued that there is no discontinuity in the problem-size effect between the very small addition problems (operands 1 to 4, assumed to be solved through compacted counting procedures) and other (medium) small problems (in which fact retrieval should occur). Instead, they observed a general performance difference between *n* + 1 and *n* + *m* problems in small additions. In addition, they showed that the problem-size effect in very small additions can also be explained through properties of an arithmetic fact network (the network interference model; Campbell, 1995). Chen and Campbell also reported evidence from other studies using the operator priming task demonstrating facilitation effects in both additions and multiplications (Chen & Campbell, 2015a) and even in zero- and one-problems (e.g., *n* + 0, *n* × 1; Chen & Campbell, 2015b). Finally, they emphasized that the assumption that adults have automatized the inefficient sum-counting strategy (i.e., count all operands) seems unlikely given that children typically abandon this strategy rather early in their development and replace it with a more efficient min-counting strategy (i.e., start counting from the larger operand; see also Chen, Loehr, & Campbell, 2019).

Because the compacted-procedure account assumes that the (counting-like) procedure is unconscious and very fast, it can hardly be distinguished from fact retrieval with behavioral measures, in particular with reaction times and strategy self-reports. Adding the neurophysiological level of analysis may provide crucial insights into these arithmetic problem-solving processes in order to advance the debate (De Smedt & Grabner, 2015). In particular, EEG correlates may help to answer the critical question of whether similar or distinct cognitive processes occur while solving very small single-digit additions and multiplications. Due to its high temporal resolution, the individual problem-solving process in each trial (from problem onset to response) can be captured with high accuracy. In addition, EEG measures turned out to be highly sensitive to cognitive processes in arithmetic problem solving so that subtle differences between operations can be detected (e.g., Grabner & De Smedt, 2011, 2012; Tschentscher & Hauk, 2016).

To date, there are only three studies that used EEG to study arithmetic and compare single-digit multiplications and additions (Wang, Gan, Zhang, & Wang, 2018; Zhou et al., 2011, 2006). However, their results remain inconclusive regarding our question of investigation because of two major limitations. First, in all studies small and large single-digit problems were analyzed together. Therefore, it is unclear to what extent activation differences emerge in very small, medium small, and large problems. A direct test of the compacted-procedure account would require a specific focus on the very small problems (with operands from 1 to 4). Second, all studies administered a (delayed) verification task, which hinders the comparison with the results from reaction time studies (Barrouillet & Thevenot, 2013; Uittenhove et al., 2016), in which participants had to actively produce the answer.

Most recently, Wang et al. (2018) used a modified operator priming paradigm (operator presented 150 ms before the operands, verification 1500 ms after the operands) to test whether single-digit additions show a greater recruitment of an executive function network than multiplications. They focused on induced (oscillatory) EEG activity and observed larger theta power increase in additions over midline and right hemisphere areas as well as larger lower alpha phase locking in anterior and central areas. In both cases, however, the effects already emerged during priming (when the operation sign was shown) and extended only to 200– 400 ms after the operands’ onset. Therefore, these findings were interpreted to reflect an “encoding difference between the two arithmetic operations rather than calculation-related processes” (p. 87) and to be “not in support of the hypothesis that procedural strategies are implicated in single-digit addition” (p. 91).

The present study directly examined the central question of whether very small single-digit additions and multiplications are solved through similar (fact-retrieval account) or different (compacted-procedure account) cognitive processes using neurophysiological data from a group of adults. Our experimental design and material was based on the study by Uittenhove et al. (2016). During EEG recording, we administered an arithmetic production task with all (non-tie) single-digit addition and multiplication problems. After the EEG session, we assessed participants’ problem-solving strategies.

We analyzed induced EEG activity, which is related to task-related coupling and uncoupling of functional networks in the brain (Bastiaansen & Hagoort, 2003; Klimesch, Schack, & Sauseng, 2005). Induced activity in the theta (around 4–7 Hz) and two alpha bands (lower: 8–10 Hz, upper: 10–12 Hz) has turned out to be particularly sensitive to (a) the cognitive processes in arithmetic problem solving (e.g., Grabner & De Smedt, 2011, 2012; Tschentscher & Hauk, 2016), in particular fact retrieval and procedural strategies, (b) operation differences between single-digit additions and multiplications (Wang et al., 2018), and (c) the arithmetic problem-size effect (De Smedt, Grabner, & Studer, 2009a; Rütsche, Hauser, Jäncke, & Grabner, 2015). For instance, it has been shown that small problems and problems self-reported to be solved through fact retrieval elicit power increase (event-related synchronization; ERS) in the theta band, particularly in the left hemisphere. Large problems and problems associated with procedural strategy self-reports, in contrast, were reflected in bilateral power decrease (event-related desynchronization; ERD) in the (lower and upper) alpha bands (De Smedt et al., 2009; Grabner & De Smedt, 2011, 2012; Rütsche et al., 2015).

In the first EEG analysis, we examined whether we can replicate the association of induced EEG activity with different cognitive processing strategies (fact retrieval vs. procedural strategies) in the set of single-digit addition and multiplication problems. Based on previous findings (Grabner & De Smedt, 2011, 2012; Tschentscher & Hauk, 2016), we expected higher theta ERS and smaller alpha ERD (in the lower and upper alpha band) for problems that were reported to be solved through fact retrieval compared to those reported to be solved through procedural strategies.

In the main analysis, we used Bayesian statistical methods to compare the strength of evidence for the fact-retrieval and the compacted-procedure accounts in the very small problems (operands between 1 and 4), as defined by Uittenhove et al. (2016). According to the compacted-procedure account, different cognitive processes take place during the solution of addition (i.e. procedure) compared to multiplication problems (i.e. retrieval). As a result, there should be stronger evidence for *differences* than for similarity in induced theta and alpha EEG activity. According to the fact-retrieval account, very small problems in both addition and multiplication are expected to be solved via memory retrieval and, therefore, the evidence should be stronger for *similarity* rather than for differences. Similar to Uittenhove et al. (2016), we focused on self-reported retrieved problems and conducted the analysis for all 12 very small problems, and separately, for the six very small problems without the *n* + 1 and *n* × 1 problems. Finally, we compared EEG activity between the six *n* + 1 and the other six *n* + *m* very small additions. In this analysis, we explored potential neurophysiological differences between these two problem categories, which can also be regarded as a further test of the sensitivity of the EEG data for subtle differences in cognitive processes.

## 2. MATERIAL AND METHODS

### 2.1. Participants

The sample consisted of 40 right-handed adults (20 female, 20 male) without known calculation difficulties, vision problems or psychiatric disorders. Participants were between 18 and 29 years old (*M* = 21.9 years, *SD* = 3.0). In total, 27 participants were psychology students, and the remaining 13 participants had at least a high school diploma. The study was approved by the local ethics committee. All participants gave written informed consent, and psychology students received course credit for their participation. After the end of the study, two randomly selected participants received a gift card worth € 10 each.

### 2.2. Materials

#### 2.2.1. EEG session

EEG trials consisted of addition and multiplication problems with all 72 operand combinations between 1 and 9 without tie problems. We separated the addition problems into three different problem sizes based on Uittenhove et al. (2016):

- *Very small* problems (12 problems; operands 1 to 4, sum ≤ 7)
- *Medium small* problems (28 problems; remaining small problems with sum ≤ 10)
- *Large* problems (32 problems; operands 2 to 9, sum > 10)

To facilitate comparability of addition and multiplication problems, we used the same operand combinations in the multiplications to define the problem sizes. Consequently, the operands were the same but the results differed between operations.

Because the focus of the study lies on potential cognitive differences between addition and multiplication in the very small problems, each very small problem was presented nine times. The medium small and large problems were presented twice (except for the 1 + 5 and 2 + 5 problems, which were also presented nine times). This resulted in 256 trials per operation and a total of 512 trials.

Figure 1 illustrates the timing of an EEG trial. Each trial started with the presentation of a fixation point for 2 seconds, after which the problem appeared. Participants were instructed to solve the problem as accurately and quickly as possible and to speak the solution out loud. A microphone recorded the responses, from which we computed reaction times (RT) from stimulus presentation to speech onset. Oral responses were noted by the experimenter and cross-checked using the audio recordings, which we used to categorize trials into correct or incorrect. After the response, an inter-trial interval (ITI) of 1 second followed. If the participants did not answer, there was a timeout after 5 seconds. Prior to the actual EEG paradigm, participants solved 10 practice problems to familiarize themselves with the experimental task (the practice problems consisted of single-digit tie problems, which were not part of the problems in the study).

#### 2.2.2. Solution strategy session

Data collection for self-reported solution strategies took place after the EEG session, because the strategy report could have influenced the performance of the simple arithmetic problems and thus biased the results (Kirk & Ashcraft, 2001).

First, participants read an information sheet with explanations and examples for possible solution strategies in simple arithmetic (following Campbell & Xue, 2001; LeFevre et al., 1996). In the subsequent computer-based paradigm, they solved each addition and multiplication problem from the EEG session again, but this time only once. Figure 2 shows a schematic illustration of a trial for the solution strategy session. Similar to the EEG trials, each trial started with the presentation of a fixation point (2 seconds), followed by the arithmetic problem. Participants were again instructed to solve the problems as accurately and quickly as possible, but this time participants gave their answer by typing in digits using a number pad. Again, there was a timeout of 5 seconds. Afterwards, a slide with solution strategies appeared on the screen, and participants reported their solution strategy by selecting either 1 (retrieval), 2 (counting), 3 (transformation) or 4 (other strategy). The inter-trial interval (ITI) was 1 second.

**Figure 2.**
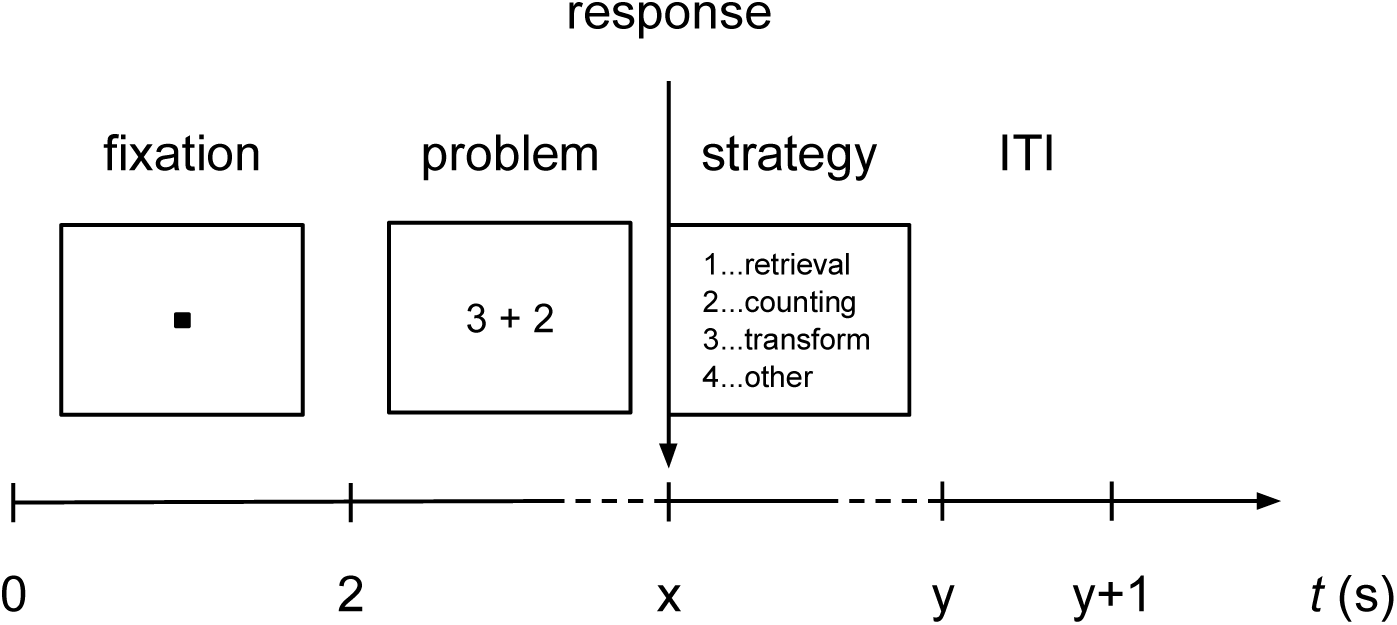
Schematic illustration of a trial in the solution strategy session.

### 2.3. Data recording

We recorded the electroencephalogram (EEG) with a BioSemi ActiveTwo system (BioSemi, Amsterdam, Netherlands) using 64 electrodes placed according to the extended 10– 20 system. Three additional electrodes recorded the electrooculogram (EOG) to detect horizontal and vertical eye movements. Two EOG electrodes were placed horizontally at the outer canthi of both eyes, and the third electrode was installed above the nasion. EEG and EOG signals were sampled at 512 Hz and lowpass-filtered at 128 Hz.

In both sessions, we implemented and presented the paradigms using the open-source stimulus presentation toolbox PsychoPy (Peirce, 2009) (http://www.psychopy.org/).

### 2.4. Procedure

In the EEG session, participants completed 512 trials in four blocks of 128 trials. Each block consisted of only one arithmetic operation, yielding two addition and two multiplication blocks. Within each block, problems were presented in a fixed pseudorandomized order. The addition and multiplication blocks were presented in alternating sequence. Half of the participants started with an addition block, and the other half started with a multiplication block. There was a short break (maximum 2 minutes) between the blocks. In the paradigm assessing self-reported solution strategies, participants completed all addition and multiplication problems without repetition. Therefore, they solved the trials in two pseudo-randomized single-operation blocks of 72 trials each. Participants started with the same operation as in the EEG paradigm. The whole test session took about 2 to 2.5 hours including mounting and de-mounting of electrodes, task instructions, and collection of self-reported solution strategies.

### 2.5. Data analysis

#### 2.5.1. Performance data

We extracted reaction times (RTs), which we defined as the time from problem onset until voice onset, by analyzing the individual recorded audio files. To this end, we used the default high frequency content onset detection algorithm (Masri, 1996) provided by the open-source aubio library (https://aubio.org/). In our RT analysis, we used only correctly solved problems, thereby discarding all trials that were either incorrectly solved or not solved at all (timeout), trials with reaction times less than or equal to 0.5 s (because these are likely false activations), and trials with technical or other problems. In total, we discarded 2.5% of all trials across participants.

#### 2.5.2. EEG data

First, we filtered the recorded signals between 0.5 Hz and 45 Hz to eliminate slow-frequency and power-line contamination. Next, we visually examined continuous activity as well as power spectral densities of all 64 EEG channels. We removed bad channels (i.e. disconnected channels or channels with excessive amount of noise) from further analyses. Subsequently, we re-referenced the remaining channels using a common average reference, and we manually marked segments containing motor activity or miscellaneous technical artifacts. Finally, we performed Extended Infomax independent component analysis (ICA) (Lee et al., 1999) to remove ocular activity from the EEG signals (Jung et al., 2000). We carried out all these preprocessing steps in Python using the open-source toolbox MNE-Python (Gramfort et al., 2013; Gramfort et al., 2014) (https://mne.tools/).

Using the clean data, we computed ERS/ERD in the theta (4–7 Hz), lower alpha (8– 10 Hz), and upper alpha (10–12 Hz) frequency bands. For ERS/ERD computation, we used only artifact-free samples from correct trials. The baseline (or reference) interval (*R*) comprised EEG data from 250 ms to 1750 ms after trial onset (during the fixation phase). The activation interval (*A*) contained the time period from problem presentation (at 2000 ms) until 125 ms before voice onset (to exclude any influence of motor and/or speech artifacts). Therefore, the activation interval contained the entire time period of arithmetic problem solving (for a similar procedure, see e.g., De Smedt et al., 2009a; Grabner & De Smedt, 2011). In contrast to fixed activation intervals, which may only capture parts of the individual problem-solving process or may include task-unrelated processes, these variable activation intervals allowed us to control for differences between individuals, task conditions, and trials.

Computing ERS/ERD values started with band power values for *R* and *A* intervals, respectively. First, we computed the medians over the respective time intervals for each trial (horizontal averaging). After that, we calculated the medians across trials belonging to the three different problem sizes (vertical averaging). This resulted in one *R* value and one *A* value per channel for each of the three frequency bands. Using these values, the amount of ERS/ERD is then equal to [(*A* − *R*) / *R*] · 100%. Positive values indicate ERS (increase in band power relative to reference interval) and negative values indicate ERD (decrease in band power relative to reference interval). Similar to previous studies (e.g., De Smedt et al., 2009a; Grabner & De Smedt, 2011, 2012), for statistical analyses, we aggregated ERS/ERD values (using the arithmetic mean) for eight regions of interest (ROIs) per hemisphere: anterio-frontal (left: FP1, AF7, AF3; right: FP2, AF4, AF8), frontal (left: F7, F5, F3, F1; right: F2, F4, F6, F8), fronto-central (left: FC5, FC3, FC1; right: FC2, FC4, FC6), central (left: C5, C3, C1; right: C2, C4, C6), centro-parietal (left: CP5, CP3, CP1; right: CP2, CP4, CP6), parietal (left: P7, P5, P3, P1; right: P2, P4, P6, P8), parieto-occipital (left: PO7, PO3, O1; right: PO4, PO8, O2) and temporal (left: FT7, T7, TP7; right: FT8, T8, TP8).

### 2.6. Statistical analysis

Operation differences in the performance data (error rates and RTs) were analyzed through traditional *t*-tests, with the effect size quantified using Cohen’s *d* separately for each problem size. Results of the self-reported solution strategies are presented descriptively for the two operations and three problem sizes.

In the EEG data analysis (ERS/ERD in three frequency bands), we performed Bayesian model comparisons using an ANOVA-like design with the factor-of-interest (strategy, operation, or problem category), hemisphere (left vs. right) and ROI (eight regions as described in the previous section). In addition, all models included the random factor participant (id) to account for repeated measures. We fitted all possible models to our ERS/ERD data, sorted the models with respect to their Bayes factors (BF), and compared the best model with a model that differs only in the inclusion/exclusion of the factor-of-interest. To quantify how much one model is preferred over the other, we computed the BF of the best model compared to the other model by dividing the BFs of the two models. The rationale behind this approach is as follows. If the model that best explains the data contains the factor-of-interest, and if comparing this model with the corresponding model without the factor-of-interest results in a large BF, the evidence is stronger for *differences* than for similarities. However, if we find that the model that best explains the data does not contain the factor-of-interest and if comparing this model with the corresponding model with this factor results in a large BF, the evidence is stronger for *similarity* than for differences.

In the first analysis, we tested whether the induced EEG activity is sensitive to retrieval and procedural strategies as has been shown in previous studies using different item sets (Grabner & De Smedt, 2011, 2012; Tschentscher & Hauk, 2016). To this end, we compared theta and alpha ERS/ERD between all (addition and multiplication) problems that were reported as fact retrieval with those reported to be solved through procedural strategies (i.e., counting and transformation) using the ANOVA factors strategy (retrieval vs. procedural), hemisphere, and ROI.

For the main EEG analysis comparing the fact-retrieval with the compacted-procedure account, a similar approach with the ANOVA factors operation (addition vs. multiplication), hemisphere, and ROI was pursued. This was done separately for all 12 very small problems and for the six very small problems without the *n* + 1 problems. Finally, we computed a similar analysis for comparing the six very small *n* + 1 and six *n* + *m* problems using the ANOVA factors problem category (*n* + 1 vs. *n* + *m*), hemisphere, and ROI.

We used the BayesFactor package (https://CRAN.R-project.org/package=BayesFactor) for the R statistical computing environment (https://www.R-project.org/) to perform these statistical analyses. Specifically, we applied the function “anovaBF” to conduct our Bayesian model comparisons. Models, priors, and methods of computation used by this function are described in Rouder, Morey, Speckman, and Province (2012).

## 3. RESULTS

### 3.1. Task performance

Table 1 summarizes error rates for addition and multiplication problems across all three problem sizes. Error rates for very small additions and multiplications were both extremely low and practically identical, and the difference between operations was not significant (*t*(39) = −0.01, *p* = .99, *d* < 0.01). Error rates for medium small additions and multiplications were also both very low and did not differ from each other (*t*(39) = 0.34, *p* = .74, *d* = 0.07). Large additions, however, were solved significantly more accurately than large multiplications, with a medium effect size (*t*(39) = −5.08, *p* < .001, *d* = −0.73).

**Table 1:** Error rates of addition and multiplication problems for all three problem sizes.

|  | Very small | Medium small | Large |
| --- | --- | --- | --- |
| **Addition** | 0.42% | 0.76% | 3.62% |
| **Multiplication** | 0.42% | 0.69% | 9.47% |

Reactions times for both operations across all three problem sizes are presented in Table 2. In all three problem sizes, additions were solved faster than multiplications. In the very small problems, the effect was negligible (*t*(39) = −2.22, *p* = .033, *d* = −0.16). In the medium small and large problems, the effects were of medium and large size (medium small: *t*(39) = −7.65, *p* < .001, *d* = −0.72; large: *t*(39) = −9.00, *p* < .001, *d* = −1.32).

**Table 1:**
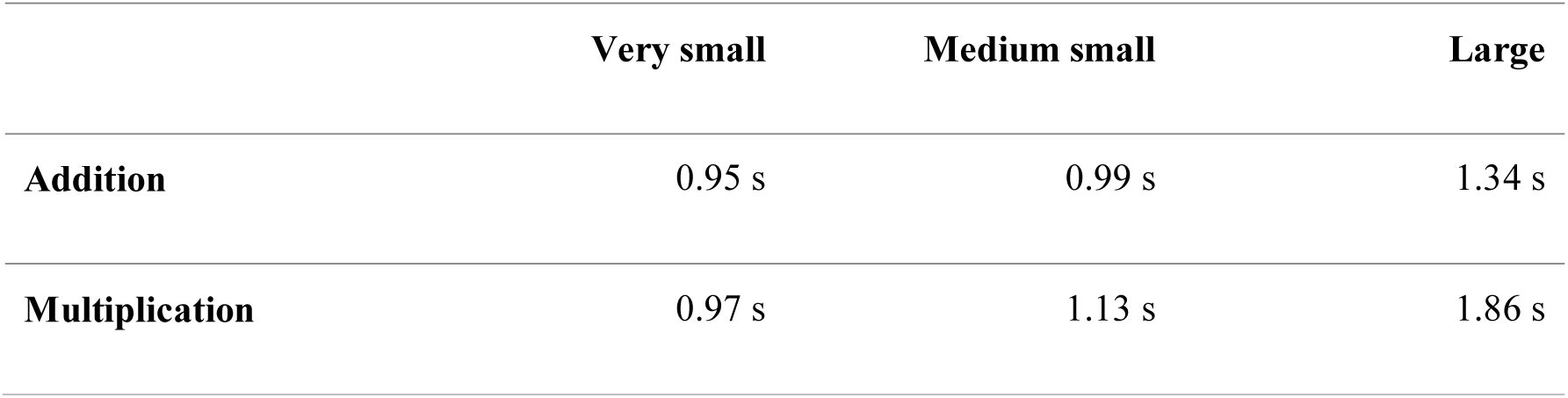
Reaction times in addition and multiplication problems for all three problem sizes.

Figure 3 illustrates the means and distributions of the RTs of all addition problems as a function of their sum. Similar to Uittenhove et al. (2016) (Figure 1, p. 294), we observed a steep increase in RTs from problems with sum 10 to those with sum 11, and a plateau for problems with sums greater than or equal to 13. In contrast to Uittenhove et al., within the small problems (sums ≤ 10), there was no systematic difference between the very small and the medium small problems but rather a monotonic increase across all small problems. Thus, based on the present RT data, a discontinuity between very small and medium small problems cannot be corroborated.

**Figure 3:**
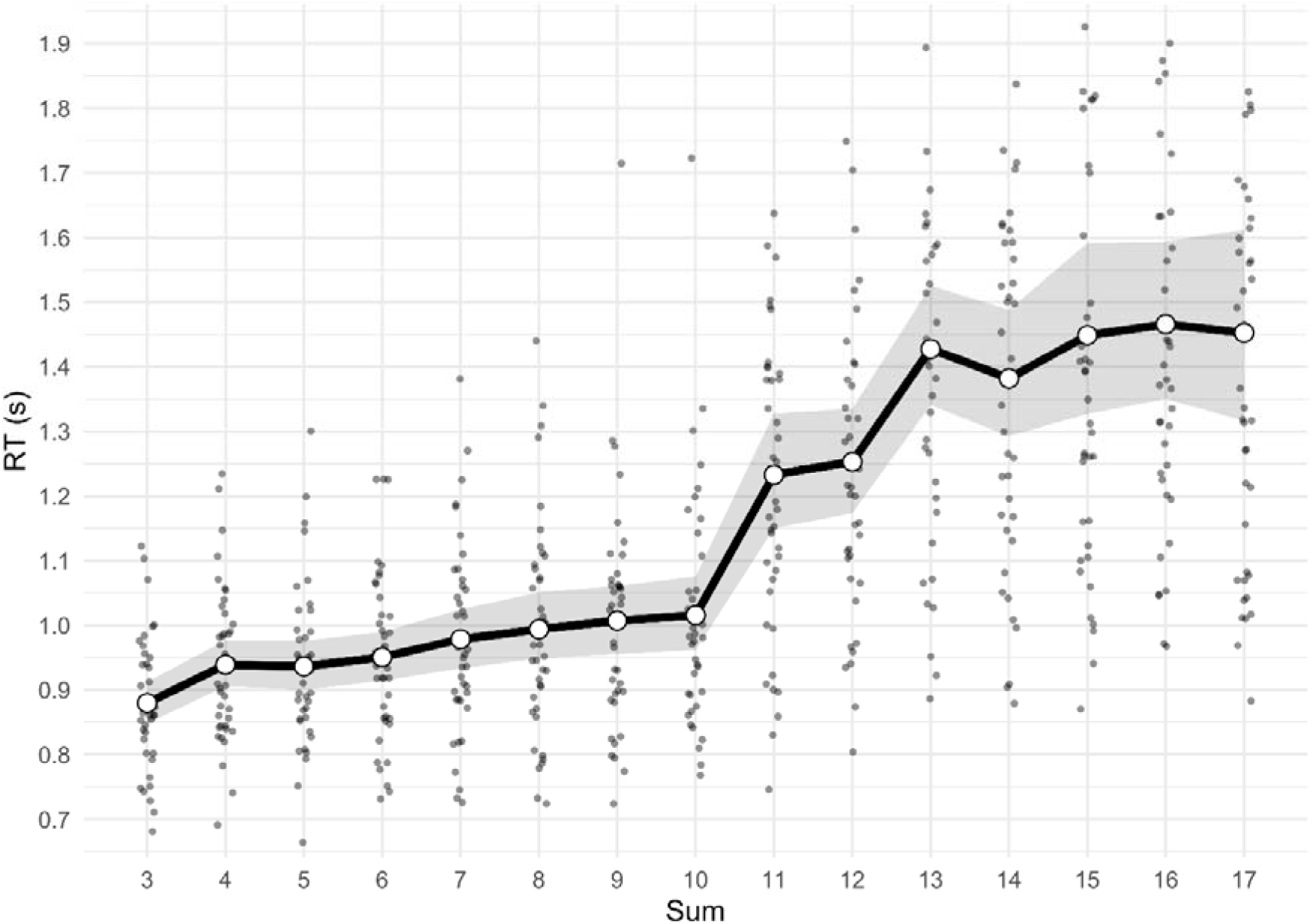
Reaction times means and distributions of all (non-tie) addition problems as a function of the sum. The gray ribbon indicates the 95% confidence interval. The y-axis is scaled from 0.65 s to 1.95 s for better visual clarity (there were 13 response times exceeding 1.95 s for sums greater than 10 that are not shown in the figure).

### 3.2. Solution strategies

The relative frequency of self-reported solution strategies is depicted in Figure 4. As expected, retrieval was the most frequently used strategy in very small (82.1% and 97.0% for addition and multiplication, respectively) and medium small problems (82.5% and 92.6%, respectively), whereas procedural strategies (counting and transformation) were used more frequently in large problems (51.3% and 36.8%, for additions and multiplications, respectively).

**Figure 4:**
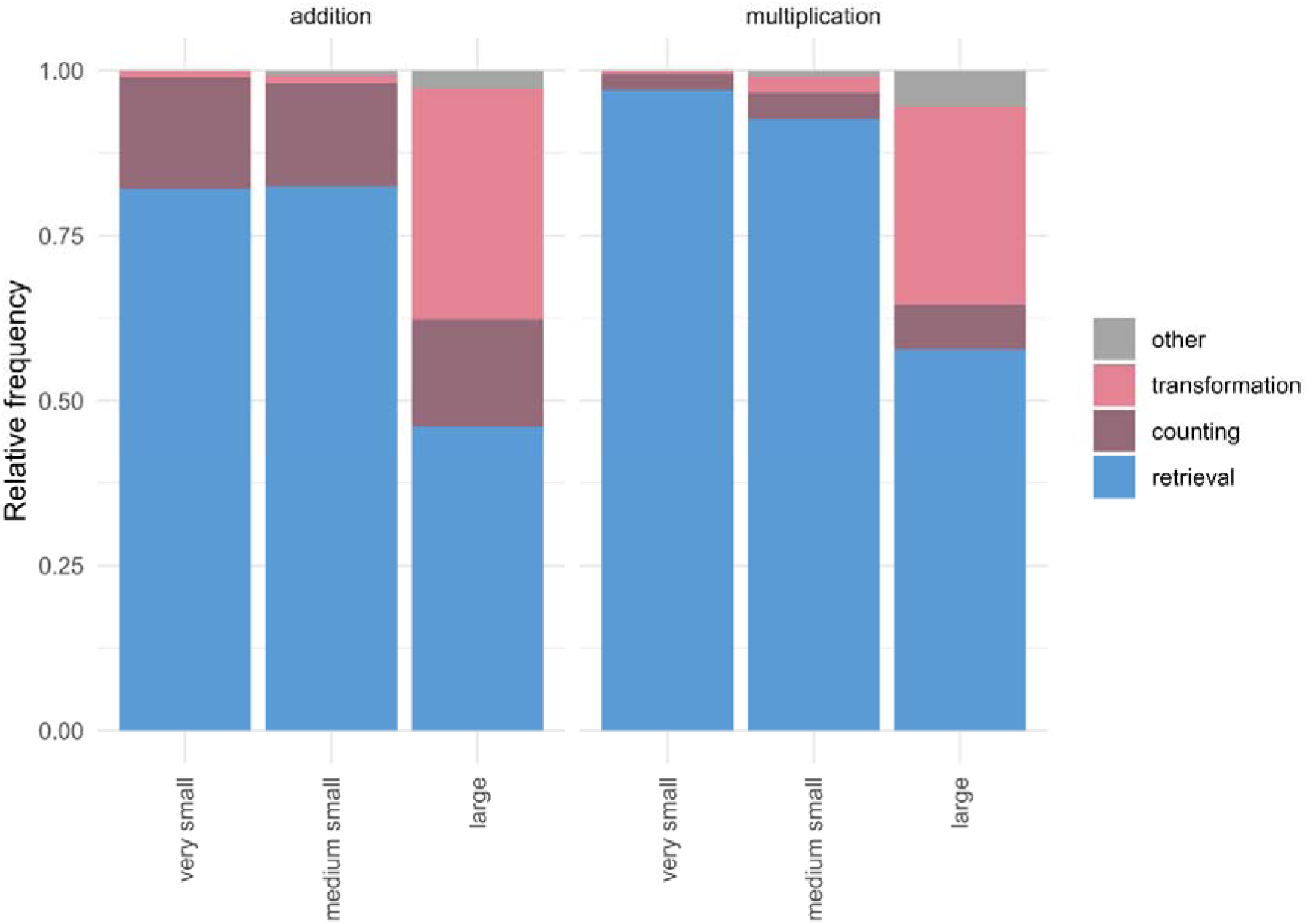
Relative frequency of self-reported strategies. Blue corresponds to retrieval, dark red and light red correspond to counting and transformation procedures, and gray corresponds to other strategies.

In order to test the sensitivity of the EEG data to self-reported solution strategies and to replicate earlier ERS/ERD studies (Grabner & De Smedt, 2011, 2012; Tschentscher & Hauk, 2016), we compared the ERS/ERD patterns between self-reported retrieval and procedural (counting and transformation) strategies. In the theta band, the best model (strategy + roi + hemi + id) contained the factor strategy and was 3.4 · 10^18^ times more likely than the model without this factor (roi + hemi + id). Figure 5 illustrates this finding and reveals higher theta ERS in several ROIs for problems reported to be solved via retrieval (retrieval problems) as compared to problems reported to be solved via procedures (procedural problems).

**Figure 5:**
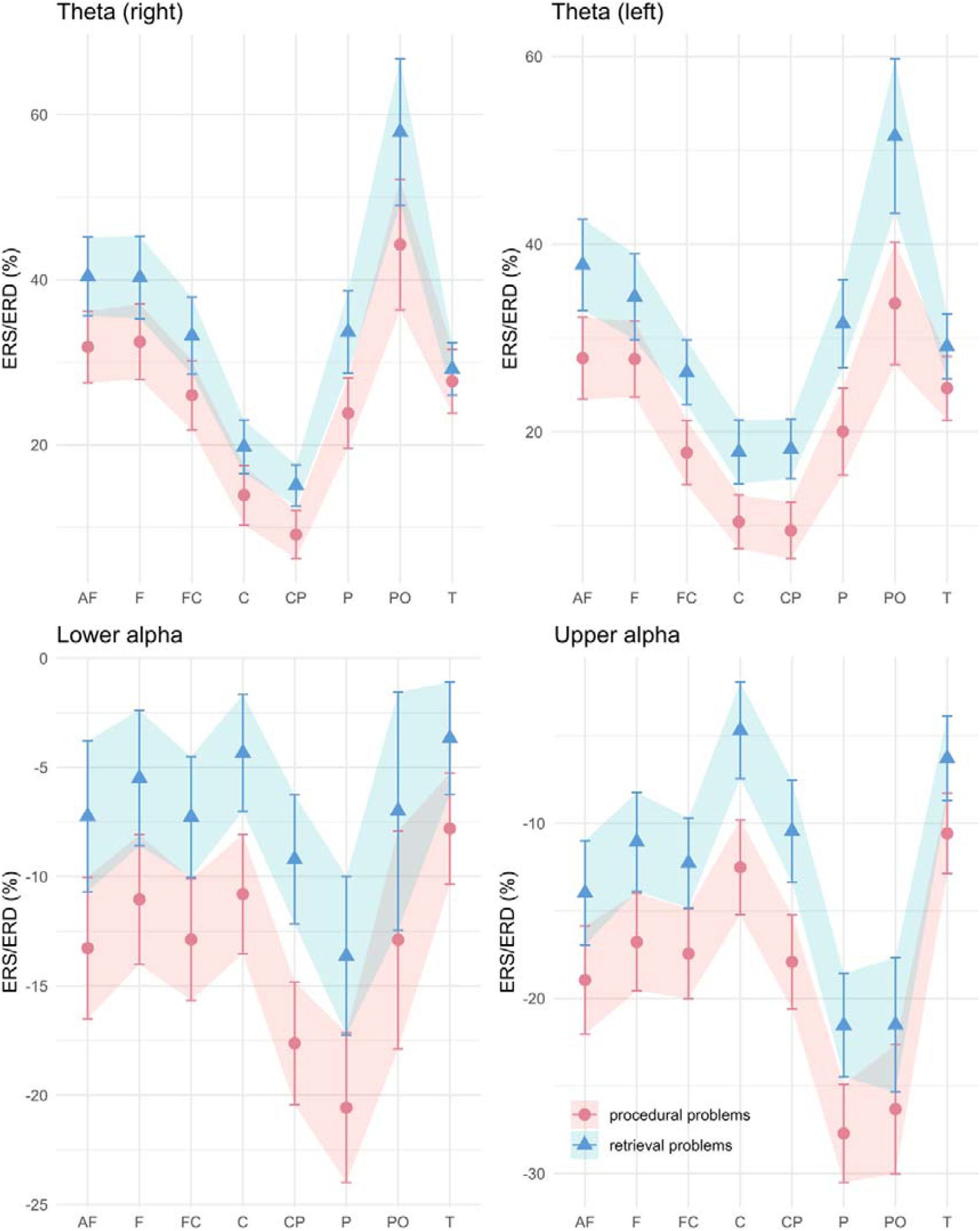
ERS/ERD in the three frequency bands (theta: upper row; lower alpha: lower row left; upper alpha: lower row right) for problems self-reported to be solved with procedures vs. fact retrieval. The ribbons indicate the standard error of the mean. ROIs are indicated on the x-axis: AF (anterio-frontal), F (frontal), FC (fronto-central), C (central), CP (centro-parietal), P (parietal), PO (parieto-occipital), T (temporal).

In the lower and upper alpha band, we also found that the best models included the factor strategy (strategy + roi + id) and were superior to the models without the factor strategy by BF of 6.0 · 10^9^ and 6.7 · 10^12^, respectively. In both bands, we observed more alpha ERD (more negative values) in procedural than retrieval problems.

Thus, in all three frequency bands, the EEG patterns strongly differed between self-reported retrieval and procedural strategies, demonstrating a high sensitivity of induced EEG activity to cognitive processes during arithmetic problem solving.

### 3.3. Analyses of retrieved very small problems

#### 3.3.1. Operation differences in very small problems

Similar to the main analysis of Uittenove et al. (2016), we first conducted the comparison between operations for all 12 very small problems in trials that were self-reported to be solved through fact retrieval. Error rates for both operations were extremely low with 0.49% and 0.46% for additions and multiplications, respectively. This difference was negligible and not significant (*t*(37) = 0.143, *p* = .89, *d* = 0.03). Reaction times were also similar with 0.96s for additions and 0.97s for multiplications. Also this difference is negligible and not significant (*t*(37) = −0.989, *p* = .33, *d* = −0.08).

The ERS/ERD for the two operations in the three frequency bands are depicted in Figure 6. For the theta band, the Bayesian model comparison resulted in a model without the factor operation (op) that best described the available data (roi + id). This model was 12.0 times more likely than the same model including the factor operation (op + roi + id). The same was true for the two alpha bands. In the lower alpha band, the best model (roi + id) was 11.6 times more likely than the same model including operation. In the upper alpha band, the BF was 6.2 in favor of the model without operation (roi + id). Thus, in all three frequency bands, there is much stronger evidence for *similarity* across operations rather than for differences, which is also reflected in the substantial overlap of the ERS/ERD in the three frequency bands (Figure 6).

**Figure 6:**
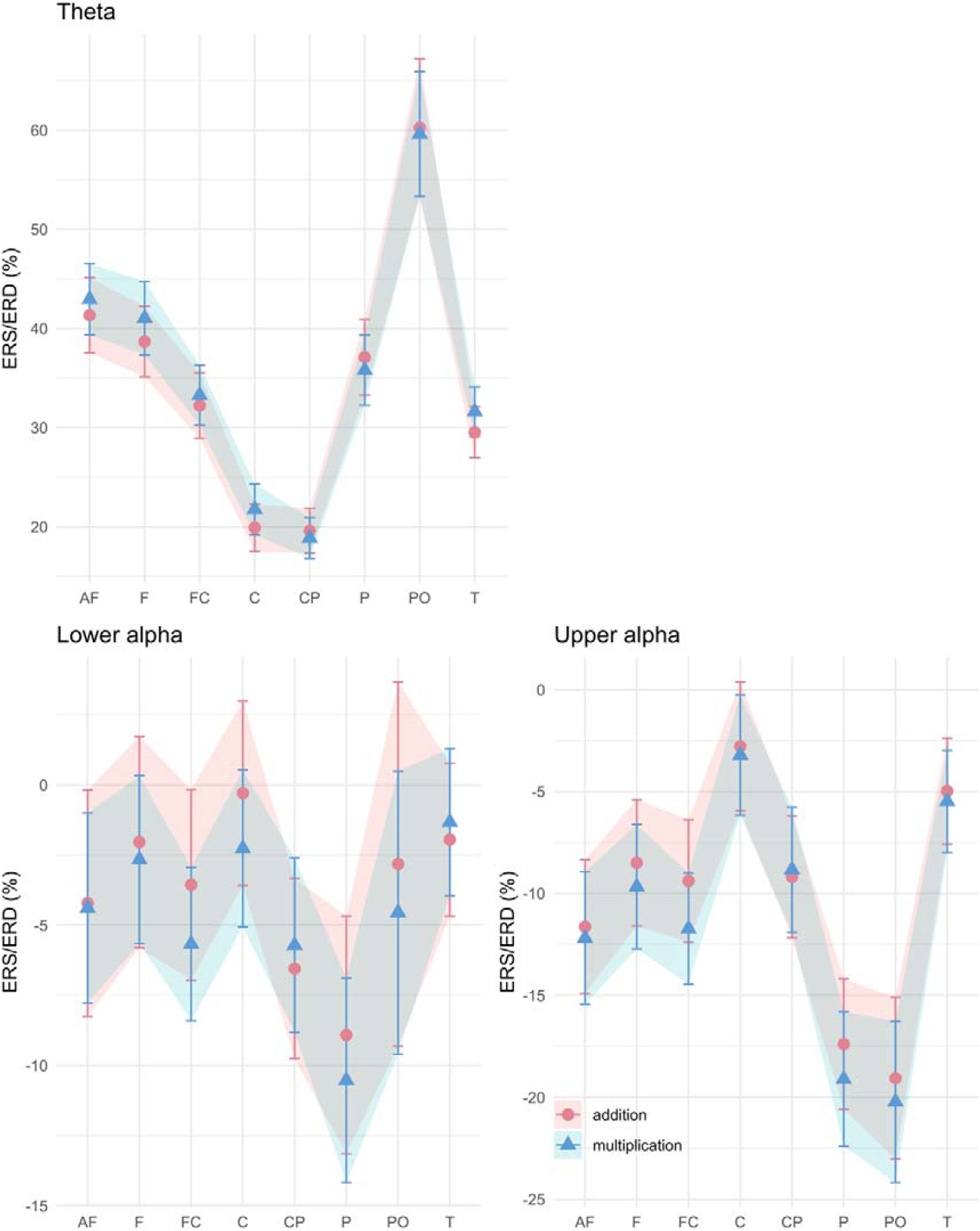
ERS/ERD in the three frequency bands (theta: upper panel; lower alpha: lower left panel; upper alpha: lower right panel) for very small retrieved addition vs. multiplication problems. The ribbons indicate the standard error of the mean. ROIs are indicated on the x-axis: AF (anterio-frontal), F (frontal), FC (fronto-central), C (central), CP (centro-parietal), P (parietal), PO (parieto-occipital), T (temporal).

#### 3.3.2. Operation differences in very small problems without n + 1 and n × 1 problems

To further corroborate our findings, again like Uittenhove et al. (2016), we conducted an additional analysis for the (retrieved) very small problems without the *n* + 1 and *n* × 1 problems, resulting in six additions and six multiplications.

Also in this problem set, error rates were similar between operations (additions: 0.69%; multiplications: 0.52%; *t*(37) = 0.45, *p* = .65, *d* = 0.11) but additions were solved faster than multiplications (0.98 s vs. 1.03 s; *t*(37) = −3.28, *p* = .002, *d* = −0.31).

Similar to the former EEG analysis, in all frequency bands, the best models only contained the ROI (roi + id) and were more likely than the corresponding model with the factor operation (op + roi + id). The resulting BF for the three frequency bands were 12.6 (theta), 15.1 (lower alpha), and 12.8 (upper alpha). These results provide further evidence for *similarity* across operations rather than for differences.

#### 3.3.3. Differences between n + 1 and n + m problems in very small additions

In a final analysis, we explored whether *n* + 1 and *n* + *m* problems within the very small problems (again only self-reported to be retrieved) differ in the accompanying EEG activity.

At the behavioral level, *n* + 1 problems were solved more accurately and faster than *n* + *m* problems (error rates: 0.15% vs. 0.69%; *t*(37) = 2.43, *p* = .02, *d* = 0.56; RT: 0.91 s vs. 0.98 s; *t*(37) = 5.31, *p* < .001, *d* = 0.46).

In the theta band, the best model contained the factor problem category (cat + roi + id) and was 4763 times more likely than the model without this factor (roi + id; see Figure 7). Similarly, in the lower alpha band, problem category was included in the best model (cat + id), which was 4.4 times more likely than the model only containing id. Only in the upper alpha band the best model solely included ROI (roi + id) and was 3.7 times more likely than the model with problem category (cat + roi + id). Even though these findings are not consistent across all three frequency bands, there seems to be stronger evidence that *n* + 1 and *n* + *m* problems in the very small additions are processed differently rather than similarly.

**Figure 7:**
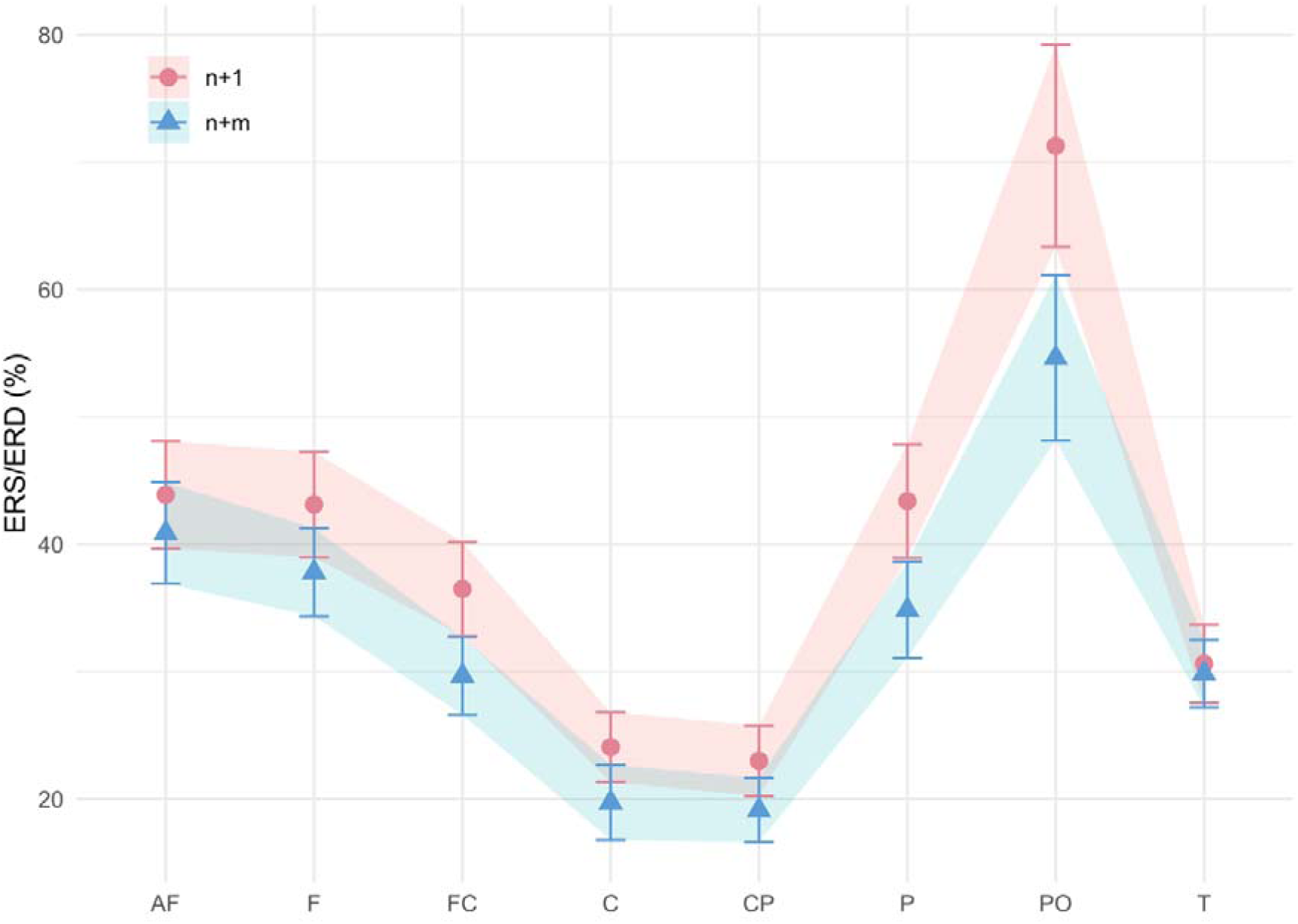
ERS/ERD in the theta frequency band for very small retrieved *n* + 1 vs. *n* + *m* addition problems. The ribbons indicate the standard error of the mean. ROIs are indicated on the x-axis: AF (anterio-frontal), F (frontal), FC (fronto-central), C (central), CP (centro-parietal), P (parietal), PO (parieto-occipital), T (temporal).

## 4. DISCUSSION

The cognitive processes underlying the solution of small single-digit additions in adults have been subject to debate in the literature on arithmetic (for recent reviews, cf. Baroody, 2018; Chen & Campbell, 2018). Two main accounts have been put forward. The *fact-retrieval account*, on the one hand, postulates that both single-digit additions and multiplications are solved through retrieval of arithmetic facts from memory (e.g., Groen & Parkman, 1972; LeFevre et al., 1996; McCloskey et al., 1991). The *compacted-procedure account*, on the other hand, hypothesizes that some single-digit additions (i.e., very small problems) are solved through a rapid and unconscious arithmetic procedure, whereas single-digit multiplications are solved through fact retrieval (e.g., Barrouillet & Thevenot, 2013; Uittenhove et al., 2016). Thus, one account postulates similar and the other different cognitive processes in solving very small problems of both operations. In the present study, we contrasted EEG correlates of solving very small single-digit addition and multiplication problems to test both accounts using a Bayesian statistical approach that compared the evidential strength for one or the other hypothesis.

The main finding of our study consists of higher evidential strength for similar rather than different EEG activity accompanying the solution of very small addition and multiplication problems that have been reported to be solved through fact retrieval. For the set of 12 very small problems, the best statistical models describing the induced EEG changes (ERS/ERD) in all three frequency bands (theta, lower alpha, upper alpha) did not contain the factor operation (addition vs. multiplication) and were between 6.2 and 12.0 times more likely than the same models including the factor operation. The picture did not change when restricting the analysis to the six very small problems without those including an operand of 1 (*n* + 1 and *n* × 1 problems), as was done in an additional analysis of RTs by Uittenhove et al. (2016). The evidence for similarity across operations was stronger than that for differences in all three frequency bands (with BFs between 12.6 and 15.1). Thus, these neurophysiological findings are clearly in support of the fact-retrieval account, suggesting that both very small additions and multiplications are solved through fact retrieval.

The present results were obtained by applying an experimental procedure that was highly similar to that of two previous behavioral studies which provided the major line of evidence in favor of the compacted-procedure account (Barrouillet & Thevenot, 2013; Uittenhove et al., 2016). First, we administered an arithmetic production task, in which participants had to actively produce the answer. Second, we presented each of the very small problems repeatedly to increase reliability in our ERS/ERD estimates. In fact, we included nine repetitions of each very small problem instead of six repetitions in the aforementioned studies. Third, we exactly followed the definition of Barrouilet and Thevenot (2013) as well as Uittenhove et al. (2016) of very small problems (i.e., problems with operands between 1 and 4). Fourth, similar to Uittenhove et al. (2016), we computed separate analyses for very small problems with and without *n* + 1 problems. Finally, we restricted our analyses to those problems that were self-reported to be solved through fact retrieval, as the main analysis in Uittenhove et al. (2016) was focused on those participants with 100% frequency of fact retrieval in the very small problems. In this vein, the present study has also overcome the limitations of previous EEG studies (Wang et al., 2018; Zhou et al., 2011, 2006), which could not answer the question of similar or different neurophysiological processes in very small problems, because they used a different experimental design as Barrouillet and Thevenot as well as Uittenhove et al.

To assess the sensitivity of induced EEG activity in theta and alpha bands for different cognitive processes in arithmetic problem solving, we conducted two further analyses. In the first analysis, we used participants’ self-reports to divide the EEG trials into retrieval trials (i.e., trials which participants reported to have solved via fact retrieval) and procedural trials (i.e., self-reported application of procedures) and tested whether the two problem-solving strategies could be differentiated in the ERS/ERD data. Evidence was substantially stronger for differences than for similarities, with BFs over 6 · 10^9^. Self-reported retrieval problems were associated with higher theta ERS and smaller alpha ERD (in both alpha bands) than self-reported procedural problems. This result replicates previous studies (De Smedt, Grabner, & Studer, 2009b; Grabner & De Smedt, 2011, 2012) and shows that even within single-digit problems, there is a strong relationship between problem-solving strategies and ERS/ERD activity in these frequency bands. In a second analysis, we contrasted the EEG activity between the *n* + 1 and the *n* + *m* problems within the critical problem set (the very small additions) to explore whether ERS/ERD is sensitive to more subtle differences in problem-solving processes. This indeed seems to be the case. In the theta band, the BF in favor of neurophysiological differences between these two problem categories was over 4700; in the two alpha bands, in contrast, the BFs were only around 4, both for (lower alpha) and against (upper alpha) such differences. This finding extends previous oscillatory EEG studies by revealing that even in very small additions, differences in neurophysiological processes can be detected between problems with and without the operand 1. This result also challenges Uittenhove et al.’s (2016) assumption of similar cognitive processes across both problem categories in the very small additions. Rather, they are more in line with Chen and Campbell’s (2018) observation of a RT discontinuity between these two problem categories. Taken these results together, the present study corroborates a remarkable sensitivity of induced EEG activity in theta and alpha bands to cognitive processes in arithmetic problem solving.

At the behavioral level, very small additions and multiplications displayed similar error rates and RTs. The error rates did not significantly differ between operations, and in the RTs there was a negligibly small speed advantage for additions compared to multiplications. In the retrieved very small problems of the main analysis, this speed advantage even disappeared. More importantly, the analysis of the addition reaction times as a function of the operands’ sum (Figure 3) only partly resembled the results by Uittenhove et al. (2016; Fig. 1). We observed a similar result pattern for problems with sums ≥ 10 with a clear initial increase in RTs reaching a plateau in the problems with solutions between 13 and 17. In contrast to Uittenhove et al., we found no boundary between very small and medium small additions; rather, there was a linear RT increase across all small problems. This finding further indicates that there is no qualitative difference in the problem-solving processes between very small and medium small problems, which is in line with the data of Chen and Campbell (2018).

The stronger neurophysiological evidence for the fact-retrieval compared to the compacted-procedure accounts adds a new level of data to the long-standing debate on the cognitive processes involved in solving (very small) single-digit additions. In fact, the evidence that has been put forward to support the compacted-procedures account was behavioral and consisted of reaction time analyses (Barrouillet & Thevenot, 2013; Thevenot et al., 2016; Uittenhove et al., 2016) and operator priming effects (Fayol & Thevenot, 2012; Roussel et al., 2002). A special case may be studies on spatial associations of arithmetic operations, which have demonstrated (rightward) shifts of attention in additions but not (or less strongly) in multiplications at both a behavioral (Mathieu et al., 2016) and a neurophysiological (Mathieu et al., 2018) level. In the latter study, using functional magnetic resonance imaging (fMRI), Mathieu et al. found that the presentation of a “+” compared to a “×” sign is related to stronger activation increases in three brain regions supporting the orienting of spatial attention. Even though the findings of distinct spatial associations in addition and multiplication are compatible with the compacted-procedure account, the authors state that this does not rule out the involvement of retrieval processes. For instance, Mathieu et al. referred to the notion by Marghetis et al. (2014) that spatial processes may complement memory-based strategies in terms of an intuitive check to limit errors. Chen and Campbell (2018) come to a similar conclusion and add that previously observed spatial attention shifts in behavior (Mathieu et al., 2016) were not limited to very small additions but emerged in both small and large problems. Notably, this is also the case in the operator priming studies (Fayol & Thevenot, 2012; Roussel et al., 2002). Against this background, the present EEG study is the first study in which neurophysiological data were used to directly test both theoretical accounts. In this vein, it can be regarded as another example for the added value of including the neural level in the investigation of cognitive theories (e.g. De Smedt & Grabner, 2015).

In conclusion, by investigating the EEG correlates of solving single-digit additions and multiplications, the present study has revealed stronger neurophysiological evidence for the fact-retrieval compared to the compacted-procedure account. This finding suggests that very small single-digit additions are solved through similar cognitive processes as corresponding multiplications, i.e., through fact retrieval. In addition, we could replicate and extend evidence on the strong sensitivity of induced EEG activity in theta and alpha frequency bands to different cognitive processes in arithmetic problem-solving. These results add a new level of analysis to the long-standing controversy regarding the cognitive processes underlying adults’ expertise in solving extensively practiced single-digit addition problems.

## ACKNOWLEDGMENTS

We are grateful to all participants and to Dennis Wambacher for the technical realization of the experiment.

